# How to normalize metatranscriptomic count data for differential expression analysis

**DOI:** 10.1101/134650

**Authors:** Heiner Klingenberg, Peter Meinicke

**Affiliations:** Abteilung für Bioinformatik, Institut für Mikrobiologie und Genetik, Universität Göttingen, Goldschmidtstr. 1, 37077 Göttingen, Germany

## Abstract

**BACKGROUND:** Differential expression analysis on the basis of RNA-Seq count data has become a standard tool in transcriptomics. Several studies have shown that prior normalization of the data is crucial for a reliable detection of transcriptional differences. Until now it is not clear whether and how the transcriptomic approach can be used for differential expression analysis in metatranscriptomics. The potential side effects that may result from direct application of transcriptomic tools to metatranscriptomic count data have not been studied so far.

**METHODS:** We propose a model for differential expression in metatranscriptomics that explicitly accounts for variations in the taxonomic composition of transcripts across different samples. As a main consequence the correct normalization of metatranscriptomic count data requires the taxonomic separation of the data into organism-specific bins. Then the taxon-specific scaling of organism profiles yields a valid normalization and allows to recombine the scaled profiles into a metatranscriptomic count matrix. This matrix can then be analyzed with statistical tools for transcriptomic count data. For taxon-specific scaling and recombination of scaled counts we provide a simple R script.

**RESULTS:** When applying transcriptomic tools for differential expression analysis directly to metatranscriptomic data the organism-independent (global) scaling of counts implies a high risk of falsely predicted functional differences. In simulation studies we show that incorrect normalization not only tends to loose significant differences but especially can produce a large number of false positives. In contrast, taxon-specific scaling can equalize the variation of relative library sizes from different organisms and therefore shows a reliable detection of significant differences in all simulations. On real metatranscriptomic data the results from taxon-specific and global scaling can largely differ. In our study, global scaling shows a high number of extra predictions which are not supported by single transcriptome analyses. Inspection of the scaling error suggests that these extra predictions may actually correspond to artifacts of an incorrect normalization.

**CONCLUSIONS:** As in transcriptomics, a proper normalization of count data is also essential for differential expression analysis in metatranscriptomics. Our model implies a taxon-specific scaling of counts for normalization of the data. The application of taxon-specific scaling consequently removes taxonomic composition variations from functional profiles and therefore effectively prevents the risk of false predictions due to incorrect normalization.

## BACKGROUND

Metagenome analysis can provide a comprehensive view on the metabolic potential of a microbial community (Eisen, 2007; Simon and Daniel, 2009). In addition to the static functional profile of the metagenome, metatranscriptomic RNA sequencing (RNA-Seq) can highlight the multi-organism dynamics in terms of the corresponding expression profiles (Poretsky et al., 2005; Frias-Lopez et al., 2008; Gilbert et al., 2008; Urich et al., 2008). In particular, metatranscriptomics makes it possible to investigate the functional response of the community to environmental changes (Gilbert et al., 2008; Poretsky et al., 2009).

In single organism transciptome studies, differential expression analysis based on RNA-Seq data has become an established tool (Marioni et al., 2008; Trapnell et al., 2012). For the analysis, first, quality-checked sequence reads are mapped to the organisms genome for transcript identification. Then the transcript counts are compared between different experimental conditions to identify statistically significant differences. Several studies have shown that read count normalization has a great impact on the detection of significant differences (Bullard et al., 2010; Dillies et al., 2013; Lin et al., 2016). The aim of the count normalization is to make the expression levels comparable across different samples and conditions. This is an essential prerequisite for distinguishing condition-dependent differences from spurious variation of expression levels.

In metatranscriptomics, already the transcript identification step can be challenging. In many cases, RNA-Seq faces a mixture of organisms for which no reference genome sequence is available. Several strategies have been suggested: de novo transcriptome assembly combined with successive homology-based annotation (Celaj et al., 2014), the direct functional annotation of reads by classification according to some protein database (Huson et al., 2011; Nacke et al., 2014; Hesse et al., 2015) or parallel sequencing of the corresponding metagenome with successive mapping of RNA-Seq reads to assembled and annotated contigs (Mason et al., 2012; Franzosa et al., 2014; Ye and Tang, 2016). For the subsequent comparison of counts between different conditions no standard protocol exists for differential expression analysis on metatranscriptomic data. Several studies and tools apply methods that have been developed for differential expression analysis in transcriptomics to metatranscriptomic count data (McNulty et al., 2013; Martinez et al., 2016; Macklaim et al., 2013). However, the question under which conditions established models from single organism transcriptomics also apply to organism communities has not been addressed sufficiently so far.

Here we present an extended statistical model for count data from metatranscriptomic RNA-Seq experiments. Theoretical considerations as well as studies on simulated and real count data show that correct normalization of the data is crucial and in general requires an organism-specific rescaling of expression profiles. This implies that a valid differential analysis should only include data that can be attributed to single organisms. The application of differential expression analysis to mixed-species data without prior separation can be found in several metatranscriptomic studies (Nacke et al., 2014; McNulty et al., 2013; De Filippis et al., 2016) as well as in dedicated pipelines for metatranscriptome analysis (Martinez et al., 2016; Westreich et al., 2016). Our results suggest that inadequate normalization of metatranscriptomic count data always bares the risk of serious errors in differential expression analysis and should be avoided consequently.

## NORMALIZATION

To clarify our arguments for an alternative normalization of metatranscriptomic data we need to explain the statistical nature of the normalization problem. We first follow the approach of Anders and Huber (Anders and Huber, 2010) for single organism RNA-Seq count data and start with a basic model for the mean of the observed counts. The expected (mean) count 𝔼 [*Y*_*ij*_] for gene (feature) *i* and sample *j* arises from a product of the per-gene quantity 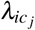 under condition *c* _*j*_ and a size factor *s* _*j*_:

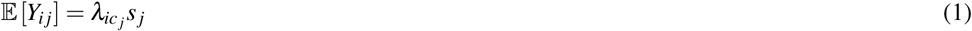

The factor 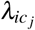 is proportional to the mean concentration of feature *i* under condition *c* _*j*_. The size factor *s* _*j*_ represents the sampling depth or library size. Usually, both factors are unknown. If we assume the *i*-th feature to be non-differentially expressed (NDE) we can represent the corresponding row of the count matrix by

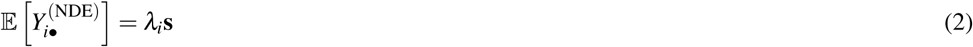

where the relative feature abundance is equal for all samples and all size factors have been comprised in the row vector **s**. Thus, for NDE features the size factors are proportional to the expected counts. If we knew which features are actually NDE, we would be able to estimate the required size (scaling) factors for normalization from the corresponding counts.

With the common choice of a mean scaling factor of 1 the scaling factors can be estimated by

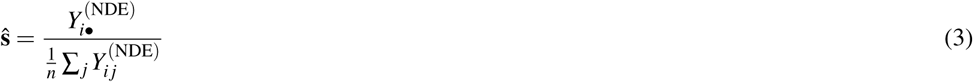

where *n* is the number of samples. The denominator in the above equation corresponds to the arithmetic mean of the counts for feature *i*. If the library sizes of the samples strongly diverge, possibly by orders of magnitude, the geometric mean is more suitable (Anders and Huber, 2010).

To make the expected counts of different samples comparable, i.e. in order to compare the feature concentrations, the columns of the data matrix are divided by the sample-specific scaling factors prior to the testing for significant differences. As above, it is common to choose an average scaling factor 1. Thus, if we would actually know an NDE feature beforehand, in principle, we could use it to estimate the scaling factors. Usually this is not the case and we need to make some assumptions. A common assumption that is used in current tools is that most of the features are NDE. Then it is possible to estimate the scaling factors by some robust statistics. In DESeq for each sample the putative scaling factors from all features are calculated and then the median of all these values is used as an estimator of the sample-specific scaling factor (Anders and Huber, 2010). The median is highly robust, with a breakdown point of 50% and therefore the estimator can be used if at least half of the data corresponds to NDE features. Without any distinction between DE and NDE features the scaling factors have also been estimated from the count sums of all samples. However, the potential shortcomings of this total count normalization have widely been discussed (Anders and Huber, 2010; Robinson and Oshlack, 2010; Soneson and Delorenzi, 2013).

In metatranscriptomics the situation is more complicated because for each organism we can have a different scaling factor. So we have to extend the above sampling model to an *N*-organism mixture that includes a matrix **S** of organism-specific scaling factors *s* _*jk*_:

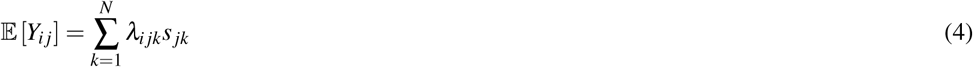

where *i*, *j*,*k* are the feature, sample and organism indices, repectively. We omitted the condition dependency (*c* _*j*_) for convenience.

In analogy to equation (2) for NDE features we have the following model for a feature row *i* of the count matrix:

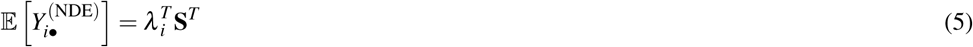

where the column vector *λ* _*i*_ contains all organism-specific rates for feature *i* and 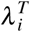 indicates transposition of this vector.

Application of the above single-organism scheme for estimation of scaling factors is only valid if the matrix of scaling factors has the following form:

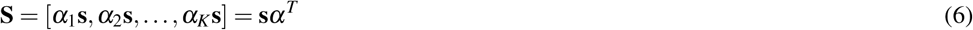

where *α* is a column vector of organism-specific abundances and **s** contains the sample-specific scaling factors, now in a column vector, which is equal for all organisms. Then we can write

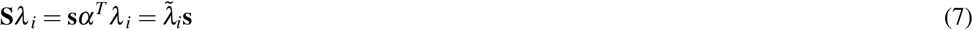

where 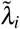 results from the dot product of the organism and the feature rates. This corresponds to equation (2) and allows to apply DESeq or other tools for single organism differential expression analysis to the metatranscriptomic count matrix. However, the underlying assumption that **S** has column rank 1, i.e. all column vectors are collinear, would be hard to justify in practice. Implicitly we would assume that the relative contributions of all organisms are constant accross all samples. In general, this assumption is not met, because for a real metatranscriptome, the organism composition of transcripts cannot be controlled and will be different for different samples. In our approach to normalization of metatranscriptomic counts we preprocess the data according to an organism-specific rescaling of seperated counts so that the recombined count data actually meet the former assumption.

## MATERIALS & METHODS

### Taxon-specific scaling and global scaling

We propose a method to prepare metatranscriptomic data for differential expression analysis. The method is referred to as taxon-specific scaling. As an essential prerequisite our approach requires that the data is first partitioned according to the contributing organisms. Then the count data matrix from each partition is normalized separately. Here, established tools from transcriptomics can be used to estimate the corresponding scaling factors. Finally, the normalized count data matrices are summed up to provide normalized metatranscriptomic count data which can be analyzed in terms of differential expression (Fig. 1). Here all statistical models and tools for count-based differential expression analysis in transcriptomics can in principle be used to identify differentially expressed features.

**Figure 1.**
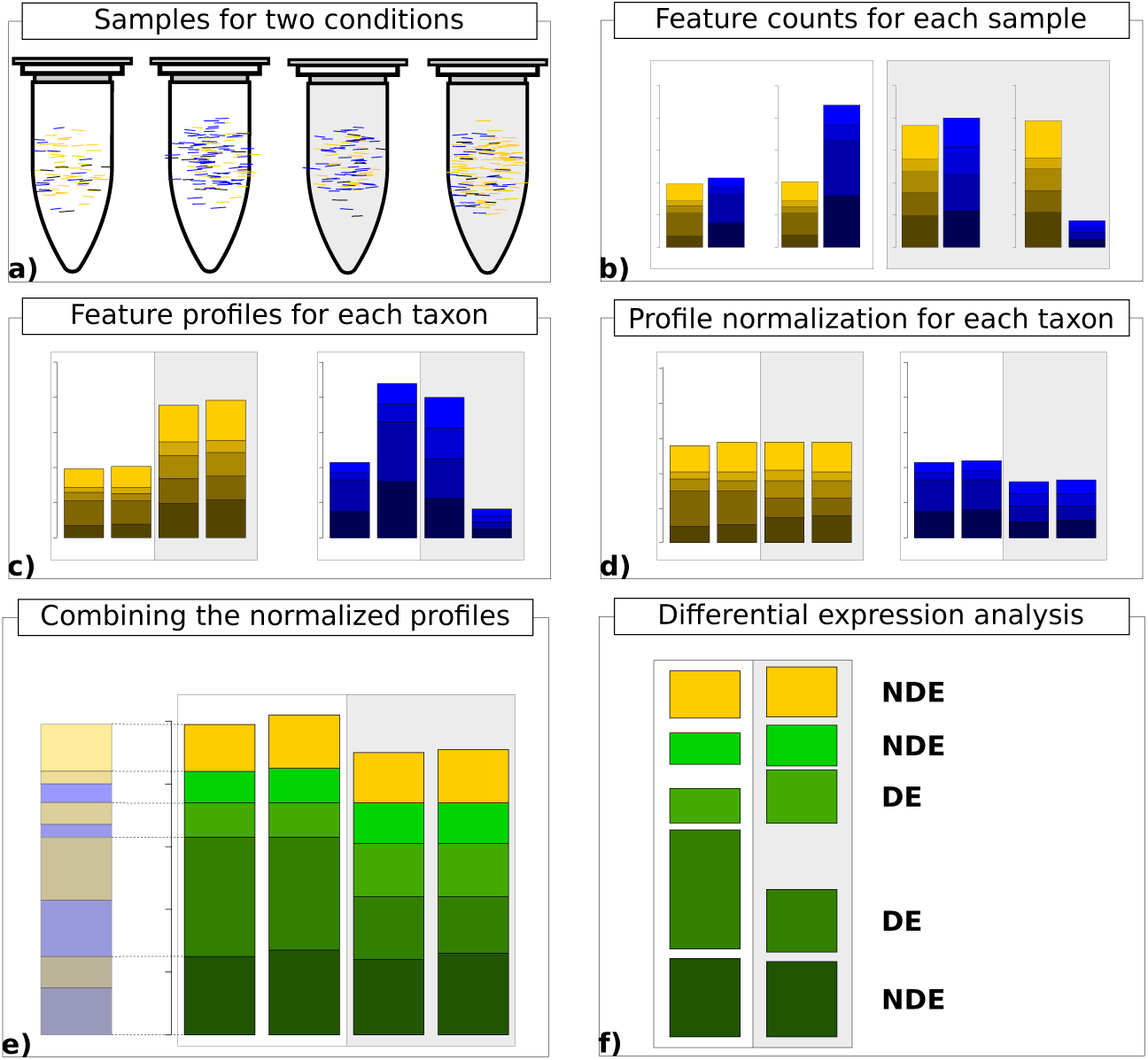
Workflow for taxon-specific normalization. a) Sequence samples from conditions A (white) and B (light gray). Assign each sequence read to taxonomic and feature categories. b) Compute feature profiles from the assignment counts. c) Obtain count matrix from taxon-specific feature profiles. d) Normalize feature profiles of each taxon-specific count matrix separately. e) Recombine normalized feature profiles of all taxa into a metatranscriptomic profile. f) Perform differential expression analysis on metatranscriptomic count matrix.

If we denote the original count matrix for organism *k* as **Y**_*k*_ and the associated vector of estimated scaling factors as 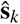 the normalized metatranscriptomic count matrix is computed by

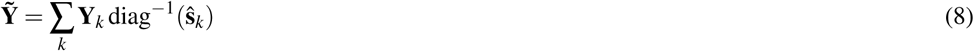

Here, the diag^−1^ operator transforms the scaling vector to a diagonal matrix with inverse scaling factors on the diagonal and zeros everywhere else. We provide an R script where we use DESeq2 for scaling factor estimation and identification of significant differences (see Additional File 1).

In principle, our method is computationally simple and the hard work has to be done beforehand in order to provide the partitioned data in terms of the organism-specific count matrices. This is the realm of binning methods and, in addition, may require sequence assembly tools to achieve a sufficient sequence length for reliable separation.

At this point, the question may arise why to get back to metatranscriptomic data when differential expression analysis could be performed for separate organisms or specific taxa. There are several reasons why the analysis of the recombined metatranscriptome data can be useful: first of all, the statistical power of organism-specific tests may be low due to decreased counts. If several organisms show the same slight difference, this difference may only become statistically significant when accumulating their normalized counts. Or a feature may show differences for single organisms but these differences may cancel out when correctly summarized. In this case the corresponding feature is not indicative for the experimental condition with regard to the whole community. Therefore the analysis of separate organism transcriptomes and the analysis of the rectified metatranscriptome data should be combined to provide a complete picture of the community response.

In our study and in the supplied R script we use DESeq2 to compute scaling factors and to identify significant differences on the basis of the normalized count matrics. We decided for DESeq2 for several reasons. It is an established tool in transcriptomics which has shown a good performance in comparative studies (Soneson and Delorenzi, 2013; Dillies et al., 2013) and which has already been used for metatranscriptome analysis (McNulty et al., 2013; Martinez et al., 2016; De Filippis et al., 2016). In particular, the estimation of scaling factors is robust and can be performed as a separate prior step apart from the computation of significant differences. The latter aspect is important for taxon-specific scaling which requires to apply the normalization independently. However, we would like to emphasize that our arguments for the taxon-specific scaling approach do not depend on a particular statistical tool and in fact the main findings of our study can be reproduced with other tools, such as edgeR (Robinson et al., 2010), SAMseq(Li and Tibshirani, 2013) or limma(Ritchie et al., 2015). In some experiments we also used edgeR and total count (TC) normalization to study the impact of different transcriptomic scaling methods.

In contrast to taxon-specific scaling, global scaling performs the normalization of metatranscriptomic data without prior separation, i.e. sample-specific scaling factors are estimated from the original metatranscriptomic counts. In general, taxon-specific and global scaling will result in distinct normalized count matrices which in turn can lead to largely differing results in differential expression analysis. We tried to show this on simulated and real count data as described in the following.

### Synthetic data generation and analysis tools

The tool compcodeR (Soneson, 2014) was used to generate all simulated data. The tool generates count data based on a negative binomial distribution model with parameters estimated from real transcriptome data (Pickrell et al., 2010; Cheung et al., 2010). If not explicitly specified, the compcodeR parameters in the R function “generate.org.mat” are used (see Additional File 1). All analyses were performed with R version 3.3.0 and DESeq2 version 1.8.2, edgeR version 3.10.5.

### Simulated metatranscriptome

A metatranscriptome arises from a mixture of various organisms, each with individual features. As a result, a metatranscriptome can include features covered by all taxa as well as features occurring only in few or a single organism. Generally, the count contributions from different organisms are not equal and vary across samples. We refer to this as the variation of the library size. Therefore, compcodeR was used to generate multiple data sets with different total count numbers to simulate the variation of organism-specific library sizes. Thereby, each generated data set mimics the contribution of a single organism. The data sets were then combined to simulate a metatranscriptomic count matrix.

As with all simulations, the data can only provide a coarse approximation of real metatranscriptomic counts which depends on particular parameters. Therefore, settings for the number of features and the number of total counts influence the results. Each organism is simulated with 100 differentially expressed features (DEF), 50 of them upregulated, and with 900 features that were non-differentially expressed (NDE).

Each data set consists of two conditions, A and B, with six samples (replicates) per condition and five organisms (Org1 to Org5) per sample. In the first three simulations, the different organism profiles are stacked, to exclude any interference between features from different organisms in the combined data. Accordingly, the final count matrix has 12 columns and 5000 rows that correspond to samples and features, respectively. The data generation process provides the necessary information to calculate the number of true positives (TP) and false positives (FP). The label *L*_*i*_ is DE or NDE according to feature *i* being differentially expressed or non-differentially expressed. The statistical test used to detect DEF, provides a p-value for each feature. The predicted label 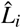 is DE if the adjusted p-value (Benjamini and Hochberg, 1995) is below a threshold of 0.05 for feature *i*. The TP and FP counts are calculated for each organism *k* individually.

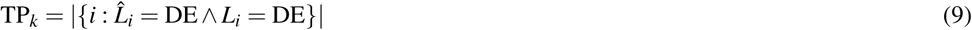

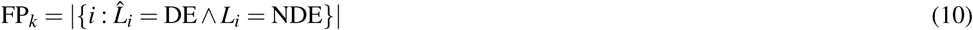

#### Simulation I: “Without library size variation”

In the first experiment we simulate the case where the libray size (LS) for each of the five organisms does not vary accross different samples. Although this is an unrealistic case we performed this simulation to verify that both normalization approaches work equally well under ideal conditions. In addition, we wanted to investigate how different organism abundances affect the identification of DEF. Each organism was assigned a fixed total count number across all samples, without variation in library size. We simulated organism Org1 with a base count of 1*e*^7^ followed by organism Org2 with 5*e*^6^, 1*e*^6^ for organism Org3 and organism Org4, Org5 with 5*e*^5^,1*e*^5^ respectively. Because data is generated without variation for the number of counts per sample, no normalization is required, i.e. the correct scaling factors for all samples and all organisms are the same (= 1).

#### Simulation II: “With library size variation”

In the second experiment we simulated a more realistic situation, with varying LS for all included organisms.

Organism base counts are identical to the first simulation but the LS is randomly increased or reduced according to a random factor between 0.5 and 2. Due to the different library sizes of the samples, a prior normalization is required.

#### Simulation III: “Condition dependent variation”

In the third simulation, we investigated to what extent a condition dependent variation of LS can affect the normalization results. Under condition A we increase LS of Org1 by a random factor between 1.5 and 2 while under condition B we decrease the LS by a random factor between 0.5 and 0.667. For Org2 the direction of change is reversed, with a random decrease under condition A and an increase for condition B. For Org1 and Org2 the same base count as for Simulation I and II is used and for Org3-5 all parameters from Simulation II are used.

#### Simulation IV: “Mixed feature effects”

In the previous simulations I-III, the generated count matrices from different organisms are stacked in the combined count matrix. By stacking the organism profiles, the provided feature labels from the data generation process can be used to validate the predictions. Now we want to analyze which effects can be observed if each feature can accumulate counts from different organisms. This case is common for real metatranscriptomic counts which arise from a mixture of organisms. Of particular interest are two effects that we refer to as ”cancellation” and ”boosting”. To simplify the analysis of these effects we restricted our simulation to a mixture of two organisms, which had the same base counts as Org1 and Org2 in Simulation I. The cancellation effect is observed when DEF that are significant in one or both organism datasets loose significance in the mixture. In contrast, the boosting effect is observed for DEF which are only significant in the combined dataset. We generated data for three samples per condition, to limit the variability of the superimposed count data. The first 100 features of the organism profiles were DE while the remaining 900 were NDE. The simulation is divided into part A, where we superimposed features with the same DE direction and part B, where the corresponding DE features of Org1 and Org2 had opposite directions.

Note that the aim of Simulation IV was not to compare the different normalization approaches but instead to demonstrate the possible effects that may result from mixed organism count data. However, the simulation cannot be used to draw conclusions about the frequencies of the effects for real data. In particular, we expect the boosting effect to be much stronger for real data where organisms with a similar response may provide correlated features that can emphasize trends or differences between conditions when superimposing their counts.

##### Part A

For this part of the simulation, we superimposed equally directed features of the two organisms. With 100 features selected as DE, the first 50 are “upregulated” followed by 50 “downregulated” features and 900 NDE features. This simulation was expected to show the boosting effect as well as the cancellation effect.

##### Part B

In part B we tried to further increase the frequency of the cancellation effect. An important aspect of identifying DEF is the difference between the mean count values of the two conditions. To bring the mean count values of the mixture for the two conditions closer together, we added the sorted ”upregulated” features of Org1 and the sorted “downregulated” features of Org2 and vice versa. The true mean values available from the data generation process are the basis for the sorting. For the generated DEF the sorting ensures that high count values in condition A from one organism are balanced with high count values in condition B from the other organism.

#### Simulation V: “False positive control”

In the final simulation, the aim was to investigate the effect of an LS shift within mixed count data without any DE features. Here we particularly wanted to measure the impact of the normalization on the false positive rate. This kind of analysis has also been proposed for transcriptomics to validate the false discovery control in differential expression analysis tools (Soneson and Delorenzi, 2013).

We used the parameters from Simulation III, but instead of six samples per condition, compcodeR generated 12 samples per condition where we only used the samples for the first condition. For Org3-5 we use the LS variation as used in Simulation III. For Org1 and Org2 the LS shift between different conditions in Simulation III is now applied between samples 1-6 (condition A) and 7-12 (condition B). The LS ranges were identical to those in Simulation III. The results of the differential expression analysis for global and taxon-specific scaling on the superimposed data were compared.

#### Error calculation of the scaling factor

The scaling error *E*_*k*_ was estimated from the difference between the sample-specific scaling factors 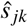 and the actual ”true” scaling factors *s*_*jk*_ for each organism *k* as provided by the simulation parameters. In both cases the factors are scaled to provide a unit mean across samples. To obtain a scaling error between 0 and 1, we compute the error by:

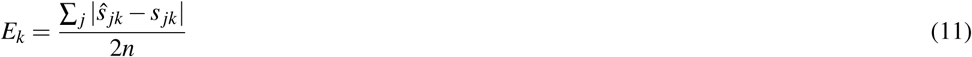

where *n* is the number of samples. In addition, we used the logarithmic measure 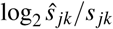 to represent the directed error.

### Metatranscriptome data

For a real data study, we chose a metatranscriptome dataset from mice gut(McNulty et al., 2013). The experiment includes 12 different species (see Additional File 2: Tab. 1) representing an artificial human gut microbial community which was inserted into germ free mice. In the original study the diet for the mice was changed at different time points. Metatranscriptomic data is available for 6 time points which provide the conditions for our analysis. The available processed count data was obtained from the European Bioinformatics Institute [http://www.ebi.ac.uk/, ArrayExpress, E-GEOD-48993] and contains gene names and the associated numbers.

Because the gene to Pfam (Finn et al., 2014) mapping is available for most organisms, we selected Pfam protein domains as features for the differential expression analysis. Each Pfam domain family is a feature in the resulting vector, including only Pfams observed at least once. We transformed the available RPKM values for the genes back to raw counts. For genes with multiple Pfam annotations, we add the raw count values of the gene to all associated Pfam features. From the available data, we constructed a count matrix for each condition and organism (Additional File 3). Here, each column constitutes a different sample and each row represents a particular feature. Because the count data from *Bacteroides cellulosilyticus WH2* did not map to gene names, all related counts are excluded from the analysis.

A differential expression analysis for all pairwise combinations of distinguished conditions was performed to compare the results of global and taxon-specific scaling. We calculated the number of DEF predicted a) with both methods with the same fold change direction, b) with both methods but with an opposite fold change direction and c) with only one scaling method. In addition, we investigated the overlap between the single organism transcriptome analyses and the differential expression analysis for the mixture. We applied a significance threshold on the adjusted p-value of 0.05 for the prediction of DEF.

## RESULTS & DISCUSSION

In the first part of our evaluation we examined the performance of taxon-specific and global scaling methods on simulated data. Because simulations I to III had been designed to provide a clear ground truth we were able to distinguish true positive predictions of DEF from falsely classified features. In the second part we show results on real metatranscriptomic count data. Here the ground truth is not known and therefore we restrict the analysis on the comparison of the results from the two normalization approaches. Because it is impossible to verify the correctness of predictions we focus on analyzing the agreement or disagreement on DEF detection in this case.

### Simulation I

In this experiment, we measured the ability to detect DEF in a metatranscriptome without variation of organism-specific libray sizes accross different samples. This situation, in principle, does not require any normalization and therefore we expected taxon-specific (”tax”) and global (”glo”) scaling to yield similar results. This is confirmed by the resulting true positive (TP) predictions of DEF for the included organisms (Fig. 2). For both approaches the number of true positives is higher for more abundant organisms due to an increased statistical power of the corresponding tests. The final profile includes 100 DE and 900 NDE features for each organism, resulting in 5000 features in total.

**Figure 2.**
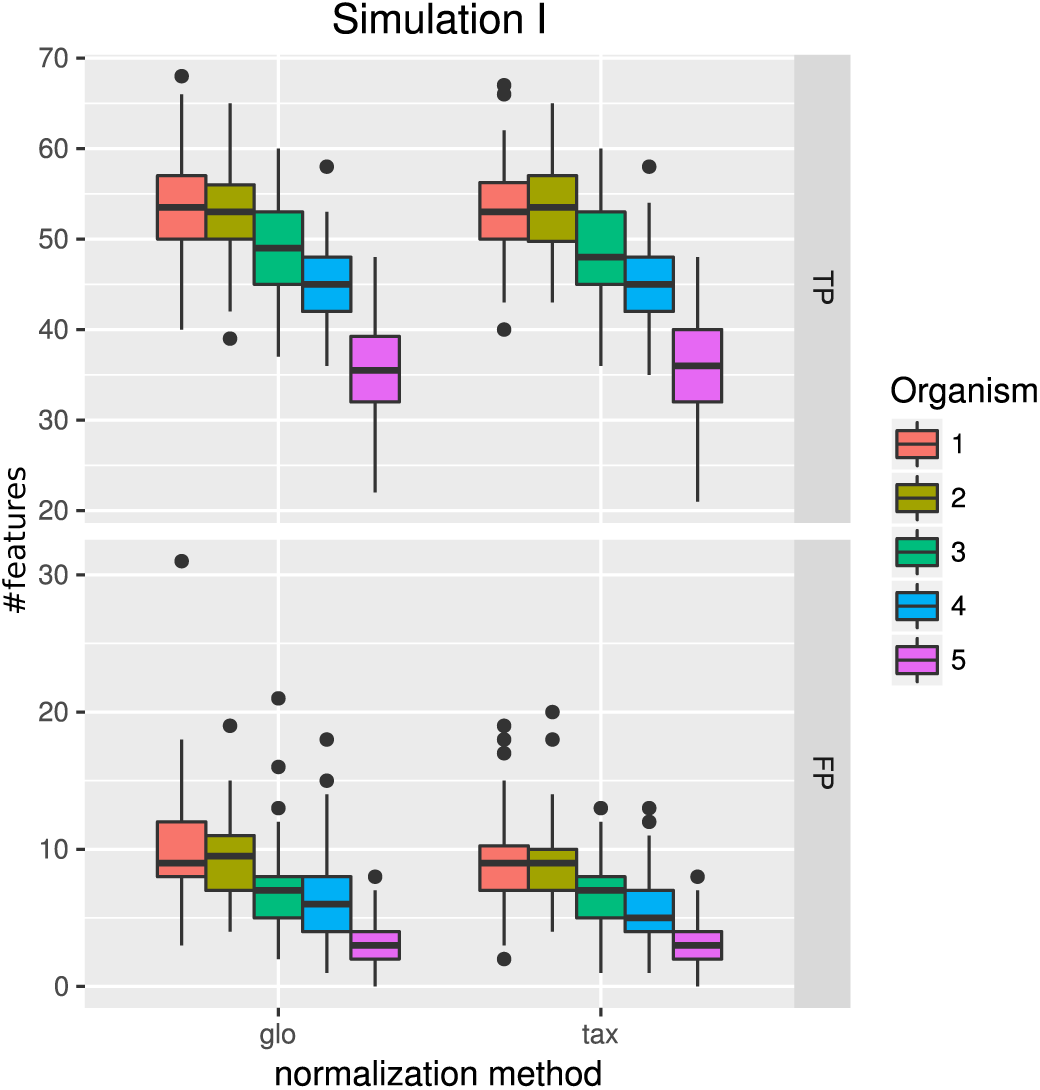
Simulation I. Number of true positive (TP) and false positive (FP) features identified with DESeq2 for global (glo) and taxon-specific (tax) scaling: Boxplots represent variation over 100 runs of the simulation.

We repeated the analysis with edgeR and TC normalizations to estimate the scaling factors (see Additional File 2: Fig. 1). For edgeR, the number of correctly identified DEF is lower for all organisms (see Additional File 2: Fig. 1). For this particular data set, the library size (LS) was correctly adjusted by DESeq2 with scaling factors close to 1 for all samples. Again both normalization approaches performed equally well. A similar picture can be expected for a varying total LS of the metatranscriptome samples as long as the relative LS of the organisms does not vary across different samples.

### Simulation II

When introducing organism-specific LS variation across samples the picture changes. For the global scaling approach the results show a decrease in the average TP rate for all organisms (see Fig. 3). This trend is also visible when edgeR and TC normalization are used for differential expression analysis (see Additional File 2: Fig 2). On the other hand, with taxon-specific scaling the results are very similar to results from Simulation I (see Fig. 2). With this method more DEF are correctly identified than with global scaling. The difference in the number of correctly identified DEF for global scaling is dependent on the parameter settings for the LS variation. For a lower amplitude of the LS variation, the TP rate for global scaling increases (Additional File 2: Fig. 3). The range of TP for the most abundant organisms Org1 and Org2 is broader with global scaling (see Fig. 3) which also shows a higher scaling error (SE) for organisms with a lower sequencing depth (see Additional File 2: Fig. 4).

**Figure 3.**
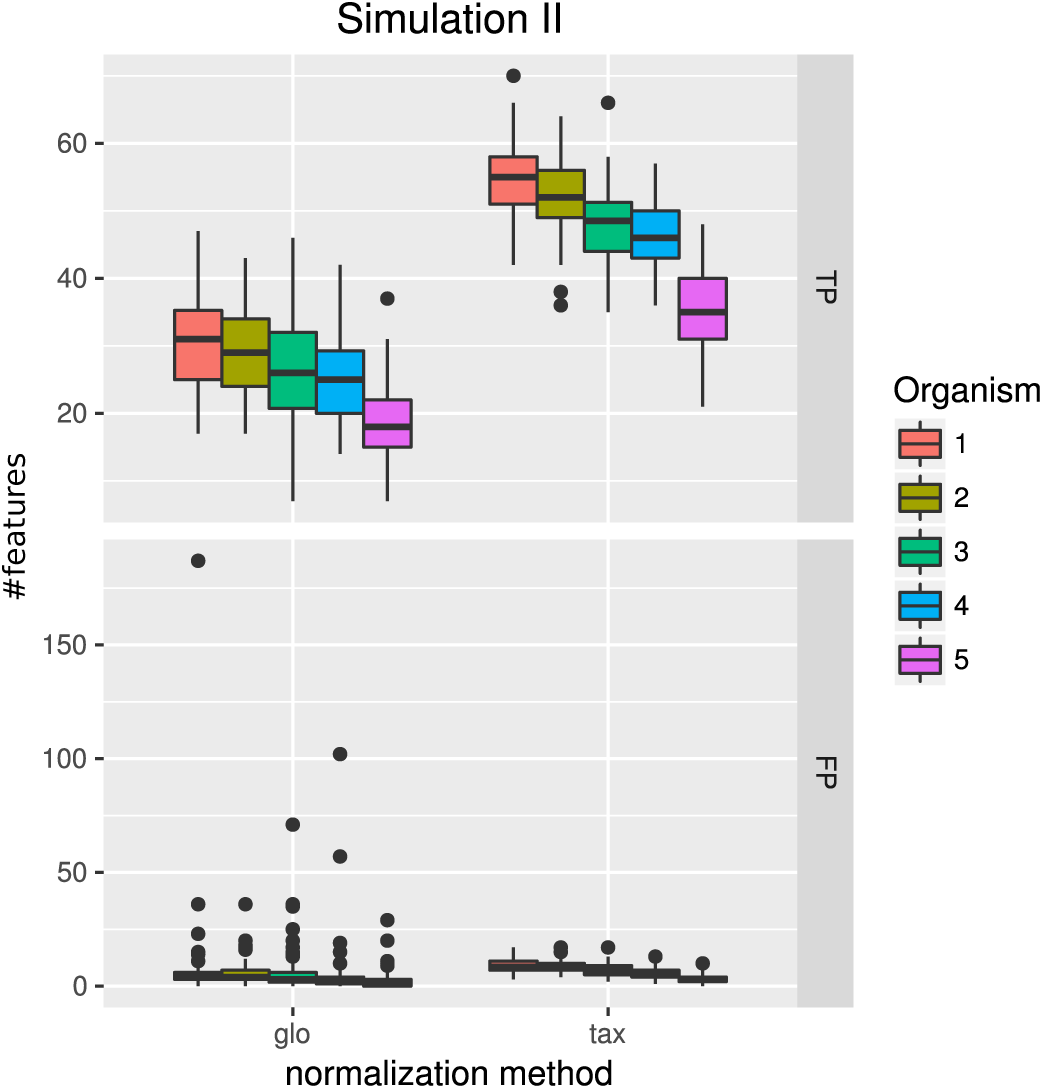
Simulation II. Number of true positive (TP) and false positive (FP) features identified with DESeq2 for global (glo) and taxon-specific (tax) scaling. FP boxplots appear compressed due to outliers. Organism order for FP is the same as for TP boxplots.

**Figure 4.**
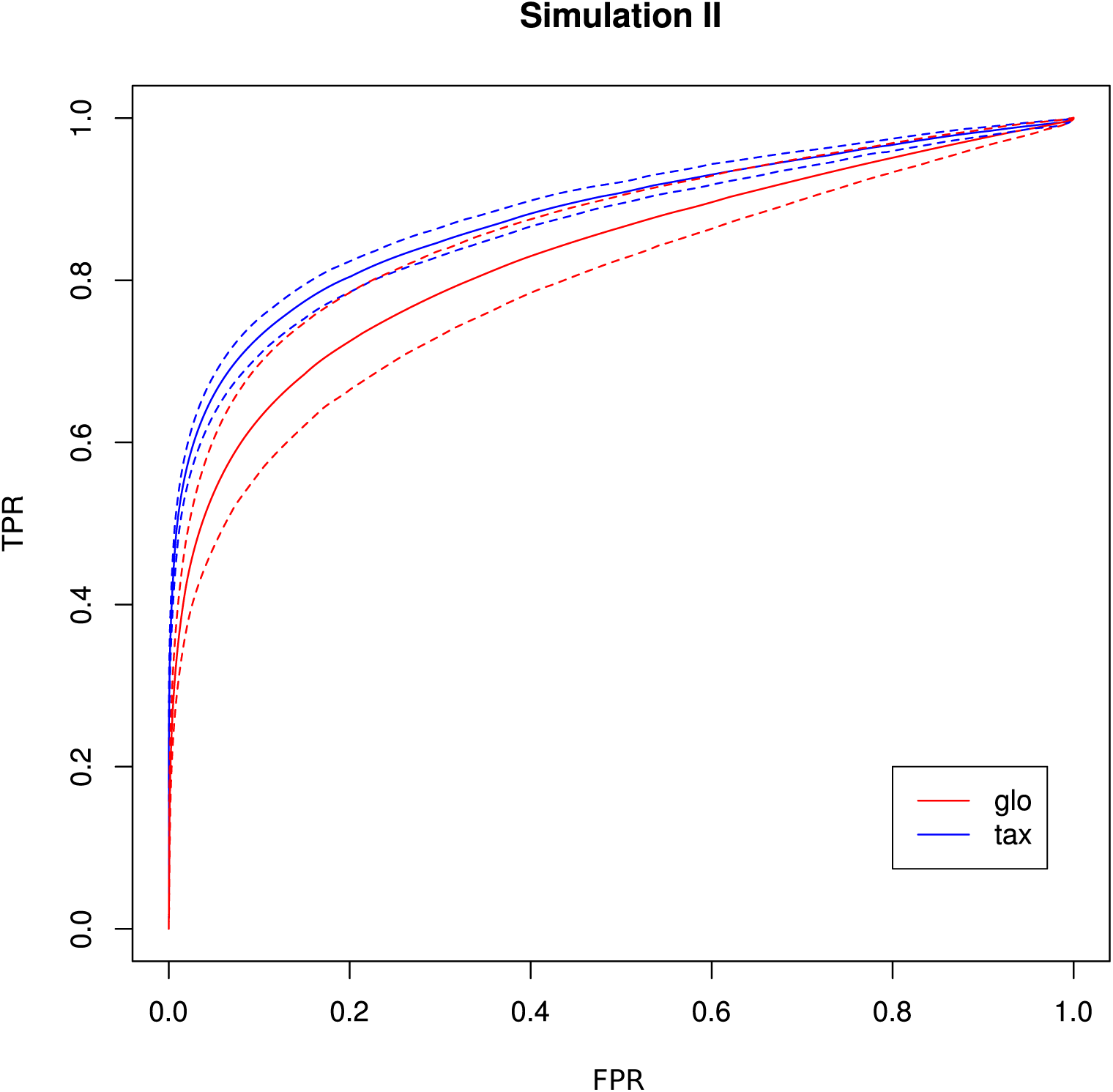
ROC curves for Simulation II. Average curve for taxon-specific scaling (blue) vs. average curve for global scaling (red) with false positive rate (FPR) on x-axis and true positive rate (TPR) on y-axis. Dotted lines above and below indicate the standard deviation for each method. The average area under curve (AUC) is 0.8776 for taxon-specific scaling and 0.8282 for global scaling.

The receiver operating characteristics (ROC) shows a higher area under curve (AUC) value for taxon-specific scaling (0.8776) than for global scaling (0.8282). The curve for global scaling also shows a higher degree of variation across different simulation runs (Fig. 4 dotted lines).

### Simulation III

With the inclusion of a condition dependent variation of the LS this simulation experiment can be viewed as a worst case study. For global scaling, the observed number of true positives is higher for all data sets compared to Simulation II (see Fig. 5).However, the number of false positive predictions explodes and even exceeds the total number of DEF (500) resulting in average TP and FP numbers of 228 (±11) and 1523 (±78), i.e.∼35 % of all features are predicted to be DEF.

**Figure 5.**
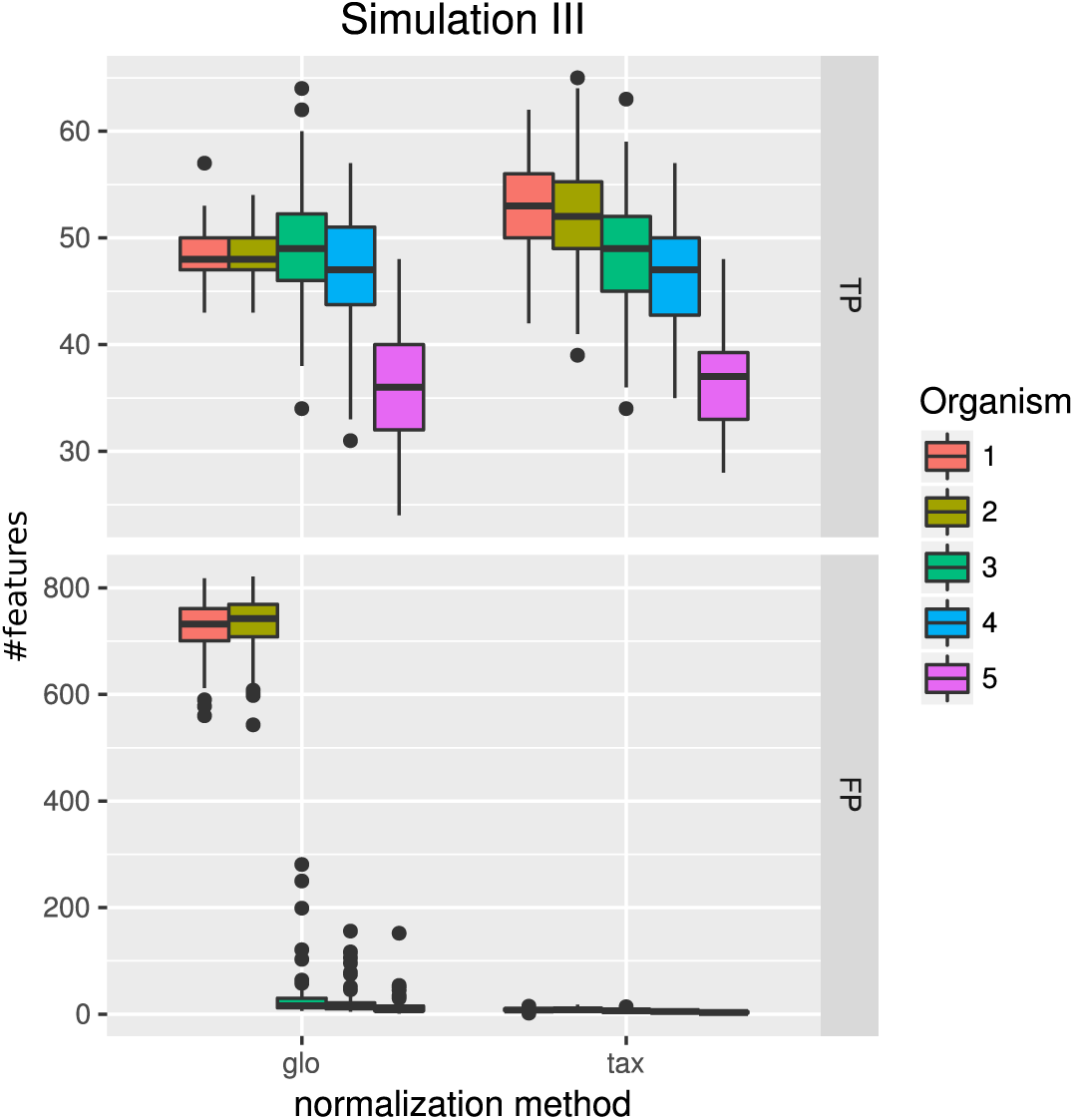
Simulation III. Number of true positive (TP) and false positive (FP) features identified with DESeq2 for global (glo) and taxon-specific (tax) scaling. Boxplots represent variation over 100 runs of the simulation.

In particular, the biggest portion of FP accumulates in features from Org1 and Org2 (see Fig. 5). Inspecting the log2 fold changes (see Fig. 6) a shift from the correct center of 0 upwards and downwards can be observed for Org1 and Org2, respectively. As a result, many DEF are identified with a wrong (opposite) direction and most of the false positive detections just reflect the direction of this shift. This situation implies a total loss of control over the false discovery rate. The results with edgeR and TC normalization show a similarly high FP rate (see Additional File 2: Fig. 5).

**Figure 6.**
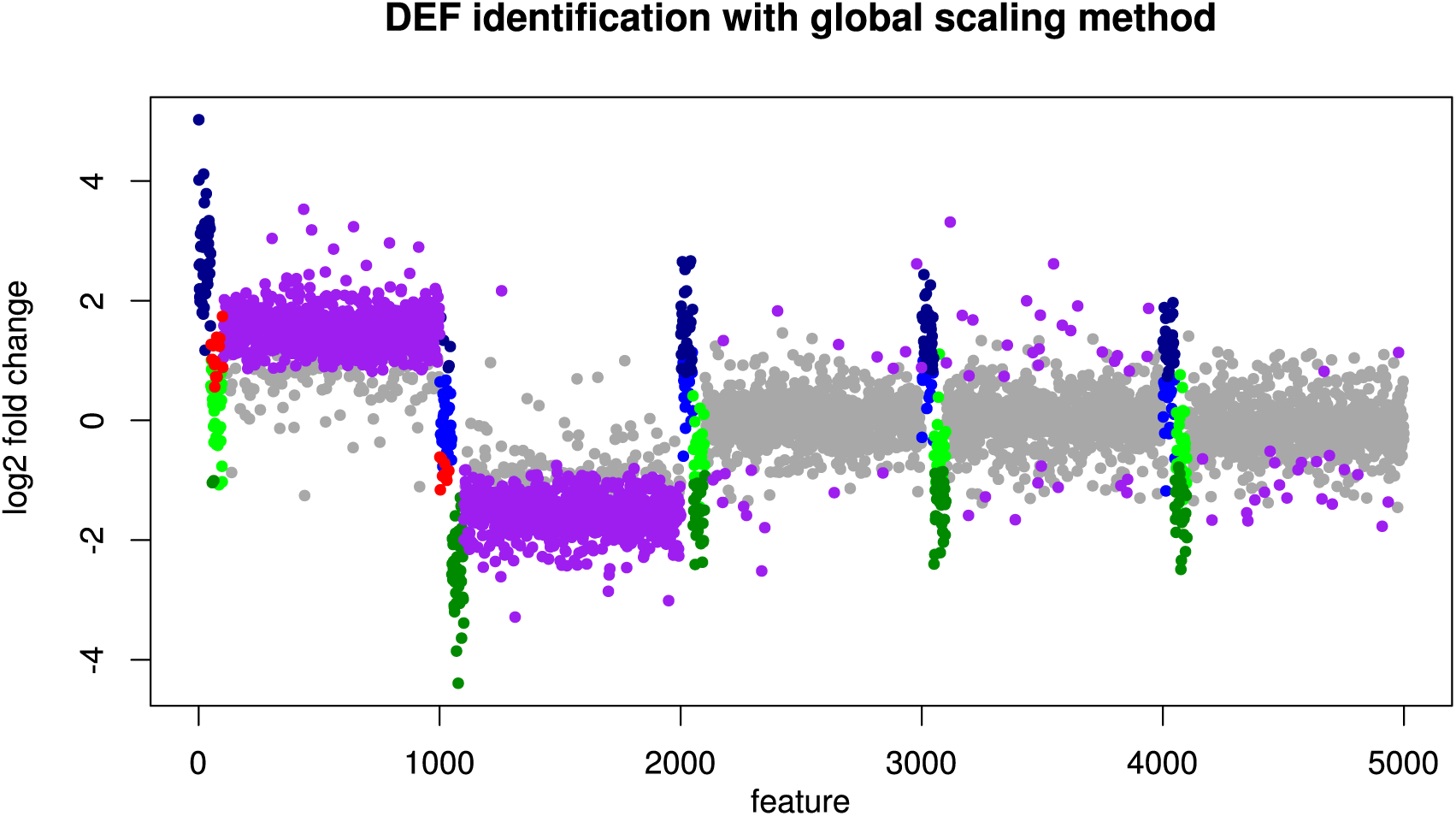
Simulation III: Log2 fold changes. Log2 fold changes of features for global normalization on one example data set. Along x-axis, features (dots) are ordered according to five stacked organism profiles, each with 1000 features of which the first 50 features are “upregulated”, and the next 50 features are “downregulated”. Gray dots correspond to correctly detected NDE features, light green dots to downregulated features which are missed and dark green dots to correctly identified downregulated DEF. Light blue dots correspond to missed upregulated DEF and dark blue dots to correctly identified upregulated DEF. Red dots mark DEF where global scaling leads to an incorrect direction. Purple dots correspond to NDE features which are incorrectly identified as significant features.

As a consequence, the ROC curve collapses for global scaling (see Fig. 7) with an AUC of 0.6396. In contrast, taxon-specific scaling does not suffer from condition-dependent LS variation and the results compare well with those of Simulation I & II showing a similar shape of the ROC curve (AUC: 0.8785). With taxon-specific scaling, the average TP across all species is 237 which corresponds to a sensitivity of ∼47 %. For global scaling the total number of predicted DEF (TP + FP) is dependent on the amplitude of the condition dependent LS shift and increases for bigger shifts.

**Figure 7.**
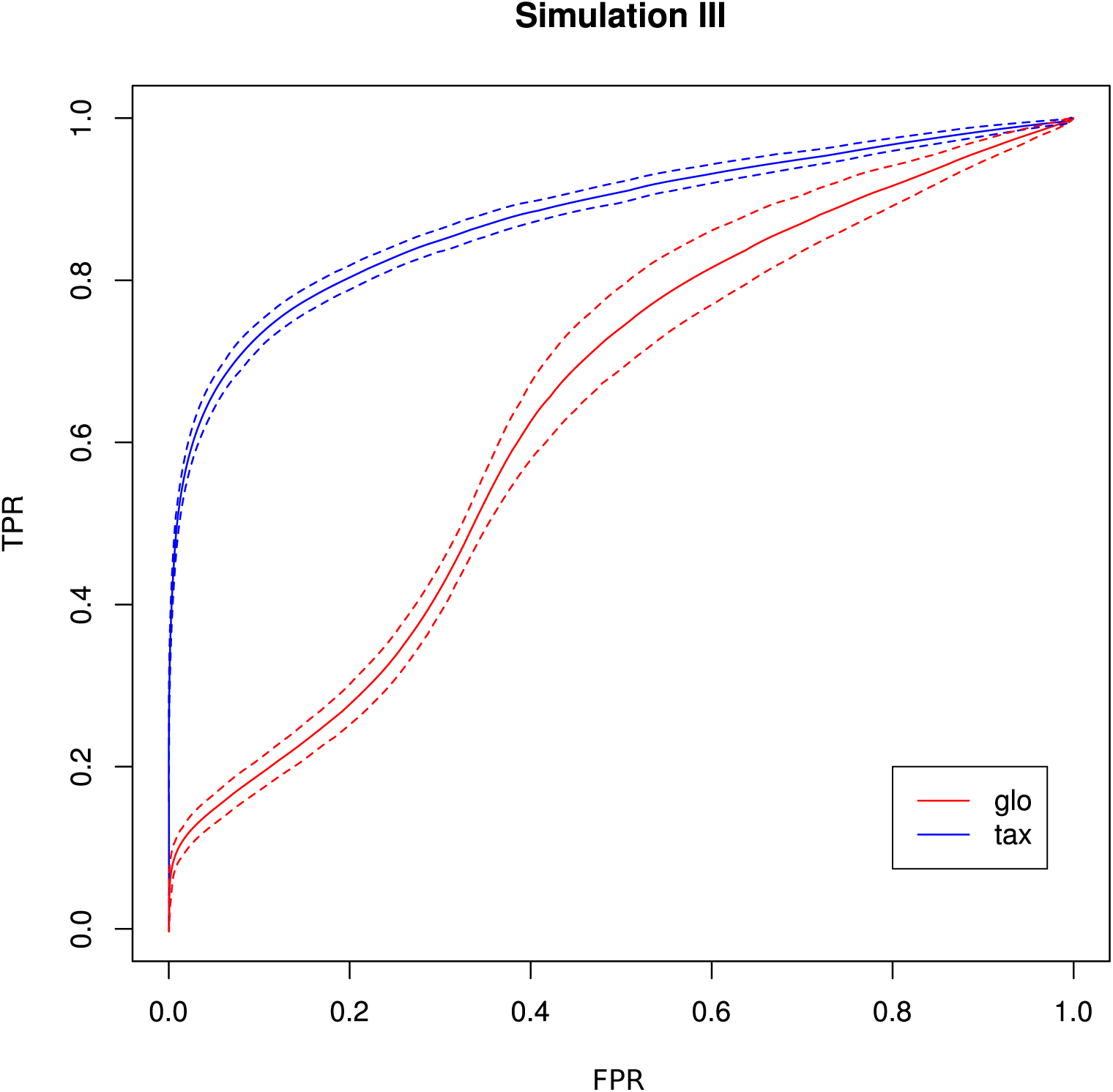
ROC curves for Simulation III. Average curve for taxon-specific scaling (blue) vs. average curve for global scaling (red) with false positive rate (FPR) on x-axis and true positive rate (TPR) on y-axis. Dotted lines above and below indicate the standard deviation for each method. The average area under curve (AUC) is 0.8785 for taxon-specific scaling and 0.6369 for global scaling.

### Simulation IV

With this simulation we wanted to analyze the effects that result from the superposition of counts from different species. In this case we do not compare the two normalization approaches. DEF that can not be detected for each organism separately may be identified as DE in the mixture (boosting effect) and DEF that can be detected for single organisms may disappear in the mixture (cancellation effect). Here we analyzed the frequencies of the different effects for the features that were labeled DE according to the data generation process. The presented numbers result from averaging over 100 iterations and for the observed effects these numbers sum up to the total number of DE-labeled features (100).

#### Part A

First, we just added the data matrices of the two simulated organisms, i.e. the upregulated and downregulated feature counts are summed up. In the results, boosting as well as cancellation effects can be observed. The chance to observe the boosting effect is low because the features of the two organisms and conditions are not correlated. Furthermore, when counts with different orders of magnitude are added, only slight changes in the mean counts can be observed. As a result, the boosting and cancellation effects only show median frequencies of 5 and 9, respectively (see Fig. 8). The main portion of features identified as DE in the mixture is also identified as DE for at least one of the organisms, followed by the fraction of features that are insignificant in all cases (see Fig. 8). Increasing the number of samples per condition from 3 to 6 further reduces the median number of boosted features (3) while also increasing the overall ability to correctly identify DE features (see Additional File 2: Fig. 6).

**Figure 8.**
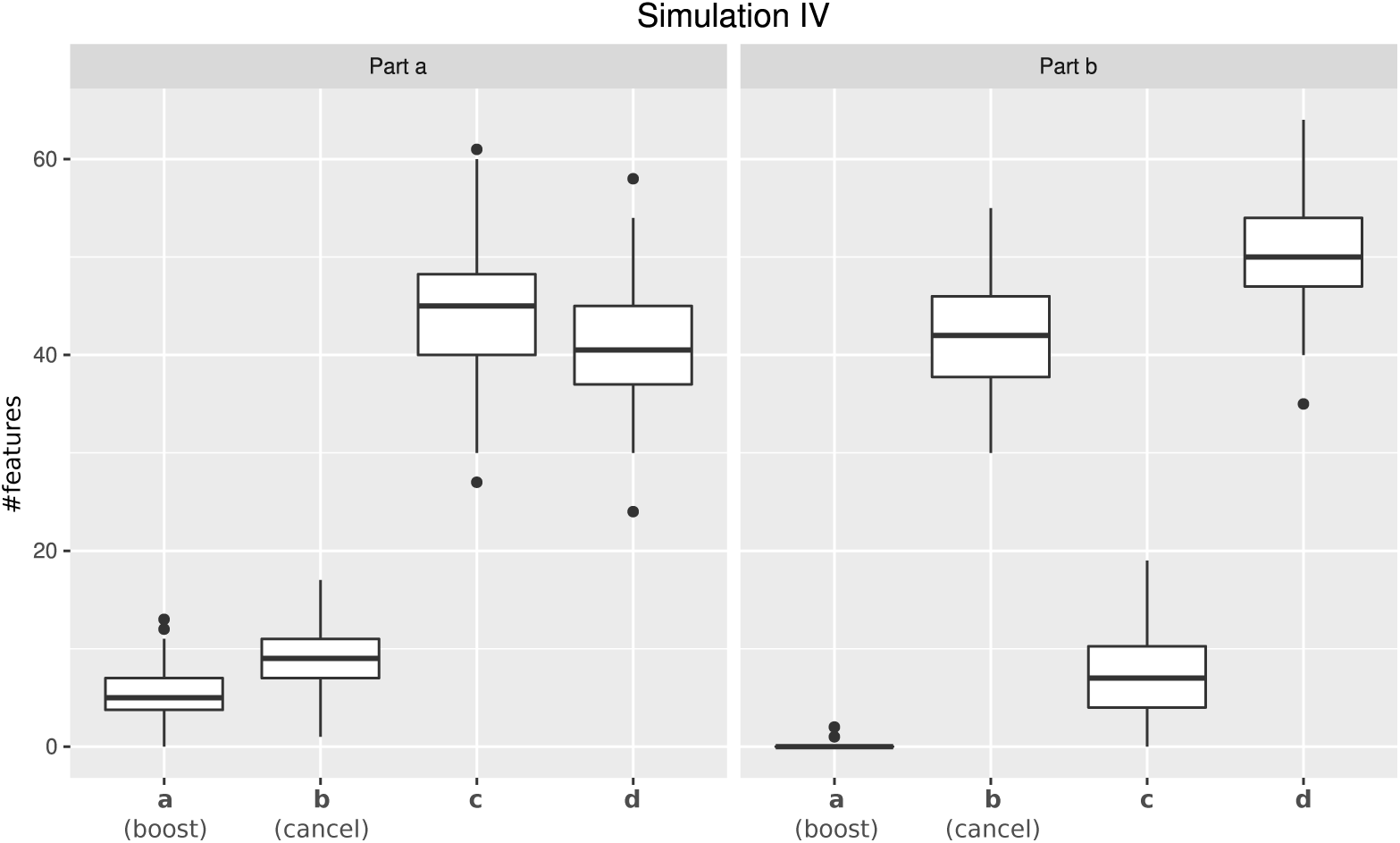
Simulation IV. Number of DEF that show a particular mixed organism effect. Effects are distinguished according to a) features not detected as DE in a single transcriptome but predicted as DE in the mixture (boosting), b) features detected as DE in at least one transcriptome but predicted as NDE in the mixture (cancellation), c) features identified as DE in at least one transcriptome and also predicted as DE in the mixture and d) features not detected as DE for both, transcriptomes and mixture. Boxplots represent variation over 100 runs of the simulation.

#### Part B

In the second part of the simulation, we changed the order of the features to intensify the cancellation effect. This was achieved by adding sorted upregulated features of one organism to sorted downregulated features of the other organism. The median number of features identified as DE for both cases, mixture and single organism analysis, is reduced to 7 and the boosting effect almost completely disappears (see Fig. 8). Due to our simulation setup, features identified as DE in one of the organisms but not identified in the mixture were the second most frequent (see Fig. 8) while features predicted as NDE for both organisms and for the mixture were most frequent.

Increasing the number of samples per condition from three to six resulted in a stronger cancellation effect and a higher number of features identified as DE for one of the organisms and in the mixture (see Additional File 2: Fig. 6). Again, the increased number of samples improves the overall detection of DEF.

### Simulation V

In the final simulation, we wanted to investigate the effect of the scaling on the false discovery rate for mixed organism count data. Therefore the data matrices (12 samples and 1000 features) for this experiment were generated without any DEF.

For taxon-specific scaling, the number of significant features is negligible with an average number of 1.6, which is well within the range of the estimated false discovery rate (FDR). In contrast, for global scaling the average number of predicted DEF is 628±22. While with taxon-specific scaling there is little if any significant difference, global scaling predicted ∼63 % of all features to be DE. Only a small number of predicted DEF results from analysis of the organism transcriptomes with a mean of 1.2, again well within the FDR range for an adjusted p-value threshold of 0.05. The results from Simulation IV show that the prevalent effect of the superposition of uncorrelated data is the cancellation effect and the boosting effect could only be observed in a few cases. However, in Simulation IV we focused on DEF that were marked as differentially expressed by the data generation process. Therefore we can exclude that the high number of DEF predicted by global scaling, results from the boosting effect.

### Real metatranscriptome data

While the objective of the simulation studies was to evaluate and compare the two normalization approaches in terms of correctly identified DEF, we do not have a ground truth for the analysis of the real data. Therefore we focused on an analysis of the (dis-)agreement in results between both approaches without assessing the actual detection performance. The analyzed data comprises Pfam counts from 11 organisms, 6 conditions according to different time points and 4 replicates per condition. An overview on predicted DEF in all pairwise condition comparisons is shown in Fig. 9.

**Figure 9.**
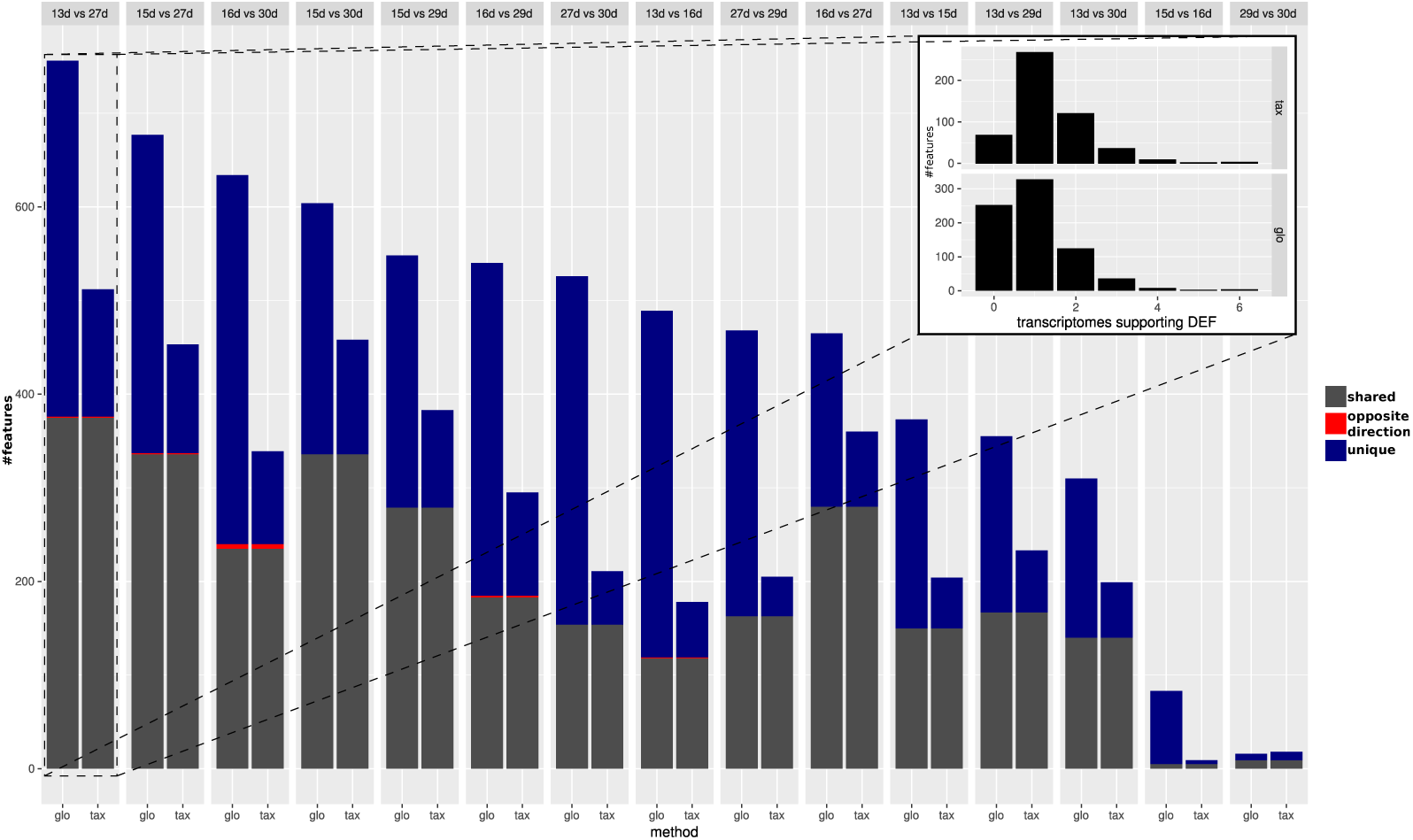
Predicted DEF for real data. Number of significant features from taxon-specific scaling (“tax”, right bar) and global scaling (“glo”, left bar) for different condition comparisons. Colors indicate shared significant features with same direction of difference (grey), shared significant features with opposite direction (red) and mutually exclusive features (purple) that are only found to be significant for one scaling method. Smaller figure: histogram for predicted DEF according to the number of single organism analyses that show a significant difference (x-axis). Upper part shows results for taxon-specific scaling and lower part for global scaling. For example, a high bar at “0” means that many features are found to be significant for the metatranscriptome which are not significant for any of the single transcriptome analyses.

For both approaches, the number of DEF peaked at “day 13” vs. “day 27” with 512 and 756 significant features for taxon-specific and global scaling. The number of DEF was low when conditions close together on the time line were compared (“day 15” vs. “day 16” or “day 29” vs. “day 30”).

For global scaling, the number of predicted DEF was generally higher than for taxon-specific scaling. For some of the comparisons, the number of extra predictions under global scaling was even higher than the number of shared predictions (see Fig. 9). The high number of extra predictions observed with global scaling is especially prevalent for the comparisons “day 15” vs “day 16” with 16 times more extra predictions than predictions shared with taxon-specific scaling and “day 13” vs “day 16” with 3 times more extra predictions than shared predictions.

We also compared the results of the mixture analysis for global scaling and taxon-specific scaling to the transcriptome analyses of the individual organisms. For 10 of the 15 comparisons, the majority of features predicted as DE with global scaling were not predicted as DE in any transcriptome (see Additional File 2: Fig. 7). In contrast, with taxon-specific scaling only the comparisons “day 29” vs “day 30” and “day 15” vs “day 16” show a higher fraction of significant features not predicted as DE in the transcriptomes. These two comparisons are also the ones with the smallest total number of predicted DEF. When taking into account the direction of the differential expression, the number of DEF predicted by both methods but with contrary regulation direction is low. Here comparison “day 16” vs “day 30” shows the highest number of significant features with an opposite direction (5).

#### Analysis of “day 13” vs “day 27”

As described in the original study, “day 13” and “day 27” each correspond to the final day of a particular diet. Because the number of predicted DEF for both scaling methods (global and taxon-specific) was the highest here, we analyze the results for this comparison in more detail.

We mapped the Pfam-annotated features which were predicted as DE for global scaling and taxon-specitic scaling to Gene Ontology (GO) terms and compared the results. With taxon-specific scaling 250 GO terms with at least one DEF mapping were identified, while global scaling resulted in 311 GO terms. GO terms associated to biological processes with a high agreement between the two methods were for example cellular amino acid metabolic process where both methods identified 7 of the 9 associated Pfams as DE and carbohydrate metabolic process with 11 DEF shared between taxon-specific scaling and global scaling. In this category, taxon-specific scaling and global scaling predicted 5 additional DEF uniquely.

GO terms with predicted DEF from taxon-specific scaling alone included magnesium ion binding and fucose metabolic process with 3 predicted DEF each. For global scaling alone, we found DNA modification, molybdopterin cofactor biosynthetic process, and RNA modification with 3, 2 and 2 predicted DEF respectively (see Additional File 4 for a complete list).

##### Extra predictions

In the condition comparison “day 13” vs “day 27”, both normalization approaches shared 376 features predicted as DE. With taxon-specific scaling, 136 extra predictions were observable while global scaling resulted in 380 extra predictions. When comparing the results of both scaling methods to the single organism transcriptome analyses, global scaling and taxon-specific scaling resulted in 252 and 69 predictions that were insignificant in all single analyses (see Fig. 9). Both methods shared an overlap of 53 DEF that were not detected in any of the transcriptome analyses. Thus, global scaling would suggest a boosting effect for ∼13 % of the features that were insignificant in all transcriptome analyses. In contrast, with taxon-specific scaling, a putative boosting effect for only ∼4% of the transcriptomic NDE features can be observed. In the single organism transcriptomes, a total of 1302 features were predicted to be DEF at least once. Both methods lead to a similar cancellation rate of ∼66% for taxon-specific scaling and ∼61% for global scaling.

The fraction of shared DEF predictions between the two scaling methods is lower if the DEF are supported by a smaller number of transcriptome analyses. For features supported by one transcriptome the agreement was ∼48%, increasing to ∼56% for two transcriptomes. In the range of three to six supporting transcriptomes, the agreement increases to ∼59%, ∼80%, 100% and 100% respectively. While the relative agreement between taxon-specific and global scaling increases, the total number of features supported by multiple transcriptomes decreases (Additional File 2: Fig. 7).

##### Scaling error

In the simulations, the differences in the estimated scaling factors for the single organism profiles in comparison to the actual scaling factors were low with a scaling error of ∼0.01. In Simulation II we found the scaling error to be high in general with global scaling (Additional File 2: Fig. 3) and in Simulation III we showed the drastic increase of features falsely identified as DE with global scaling when two organisms with condition-dependent abundance shifts were combined.

To further investigate the increased number of DEF predicted by global scaling, we compared the scaling factors estimated for the single organism profiles with the estimated scaling factors for the global normalization in the comparison “day 13” vs “day 27”. For several organisms, a pattern emerged which showed a condition specific scaling error (Fig. 10). While the scaling factors for one condition are too small, the scaling factors in the other condition are too high. As a result, global scaling leads to condition-dependent errors which may cause extra predictions of DEF because an artificial shift between the two conditions is introduced.

**Figure 10.**
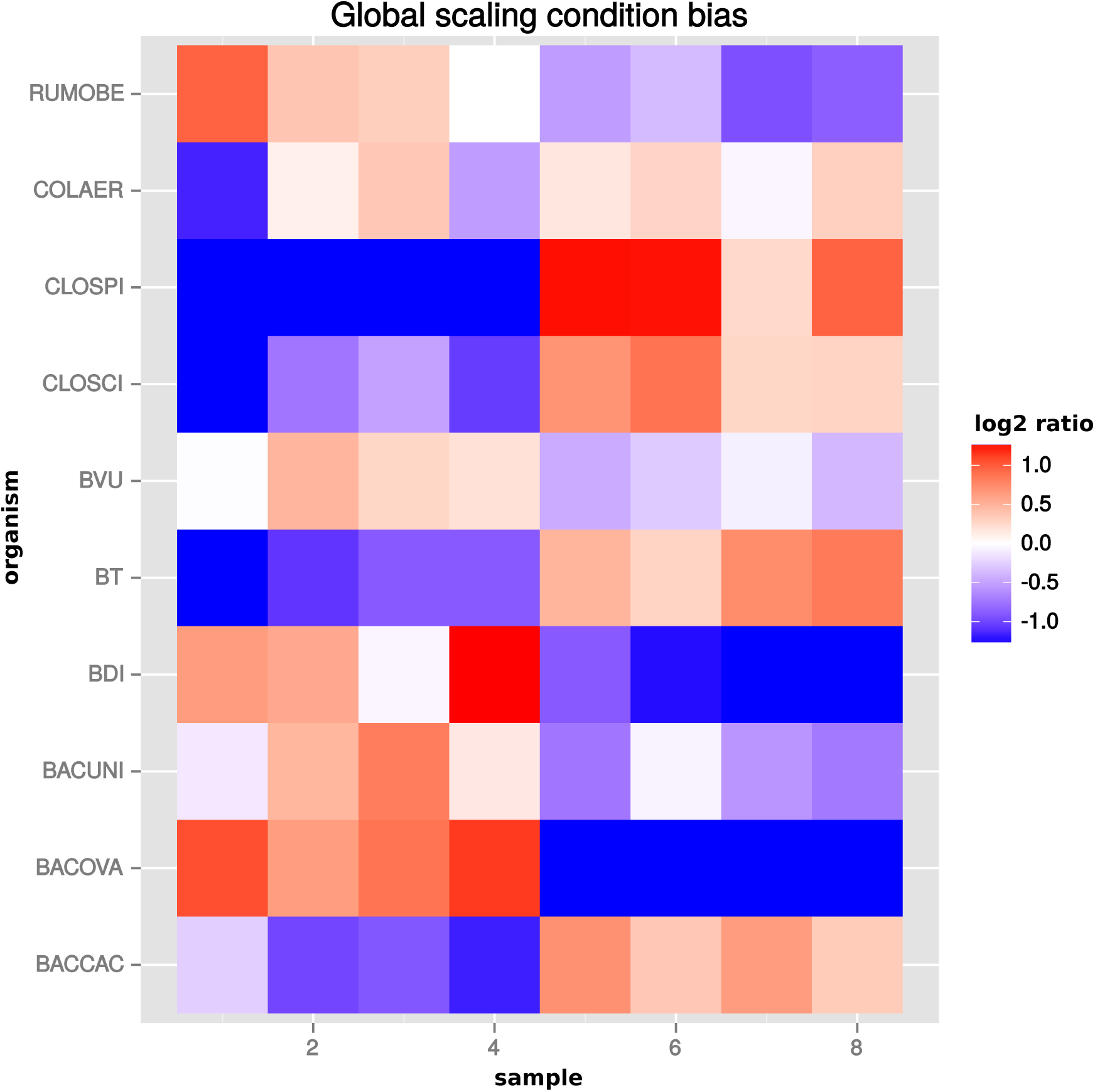
Global scaling condition bias. Direction of global scaling “error”” in terms of the log2-ratio of scaling factors from transcriptomic and global scaling. Results for different organisms in the comparison of “day 13” vs. “day 27”. For symmetry of the color range the negative log2-ratio was capped at -1.25, with error scores below that threshold showing the same color (blue). Samples 1-4 are from condition A and samples 5-8 from condition B. For the species name abbreviations see Additional File 2: Tab. 1. *D. longicatena DSM 13814* was not observed in that particular condition comparison.

Incorrect scaling is especially problematic for features, which mainly comprise counts from one organism or when counts from mixed organisms with the same scaling shift are analyzed. For a quantification we determined the number of features, for which counts from a single organism (or the mixture of organisms with the same scaling error direction) exceed 80% of the normalized counts for that feature. For extra predictions obtained only with global scaling without evidence from the transcriptome analyses, the counts from a single organism are the main contribution for 82 of 199 features. Additional 43 features are predicted from the summed counts from organisms with the same scaling shift.

##### Single feature analysis

For the comparison “day 13” vs “day 27” we now show several examples for features that reflect particular mixed organism effects or the different behavior of the scaling methods.

With regard to the scaling error discussed above, several features actually collect counts from mainly one organism (see Fig. 11, 12 and Additional File 2: Fig. 8). In addition, we found several features that show an observable boosting effect with global as well as taxon-specific scaling (e.g. Fig. 13 or Additional File 2: Fig. 9). For these features, significant differences were only observed in the combined metatranscriptome. Some of the extra predictions obtained with taxon-specific scaling are results of a putative boosting effect (see Additional File 2: Fig. 10 and Fig. 11). In contrast, some of the extra predictions resulting in a putative boosting effect were observable only with global scaling (see Fig. 11 and Additional File 2: Fig 8). In both cases, the incorrect scaling factors resulted in the detection of DE for mainly one organism. For other features, the combination of multiple incorrect scaling factors also predicted DE when global scaling was used (see Additional File 2: Fig. 12).

**Figure 11.**
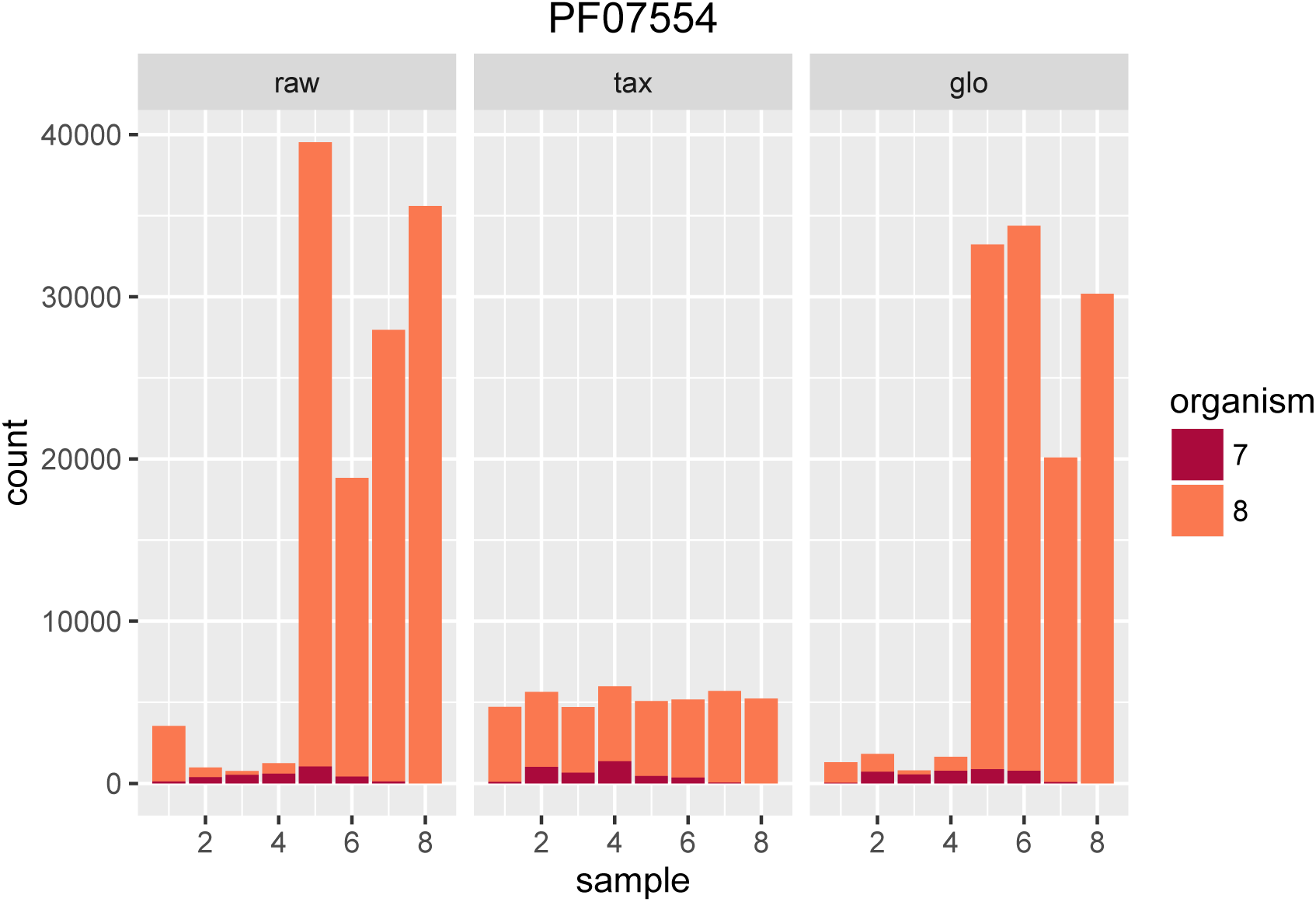
Single feature analysis (PF07554). Stacked bars in three parts (x-axis) show organism-specific counts before scaling (left), after taxon-specific scaling (middle) and after Taxon-specific and global scaling result in adjusted p-values 0.98 and 7.23^−56^, respectively. global scaling (right). Extra prediction of DEF by (mis)scaling of one organism with global scaling.

**Figure 12.**
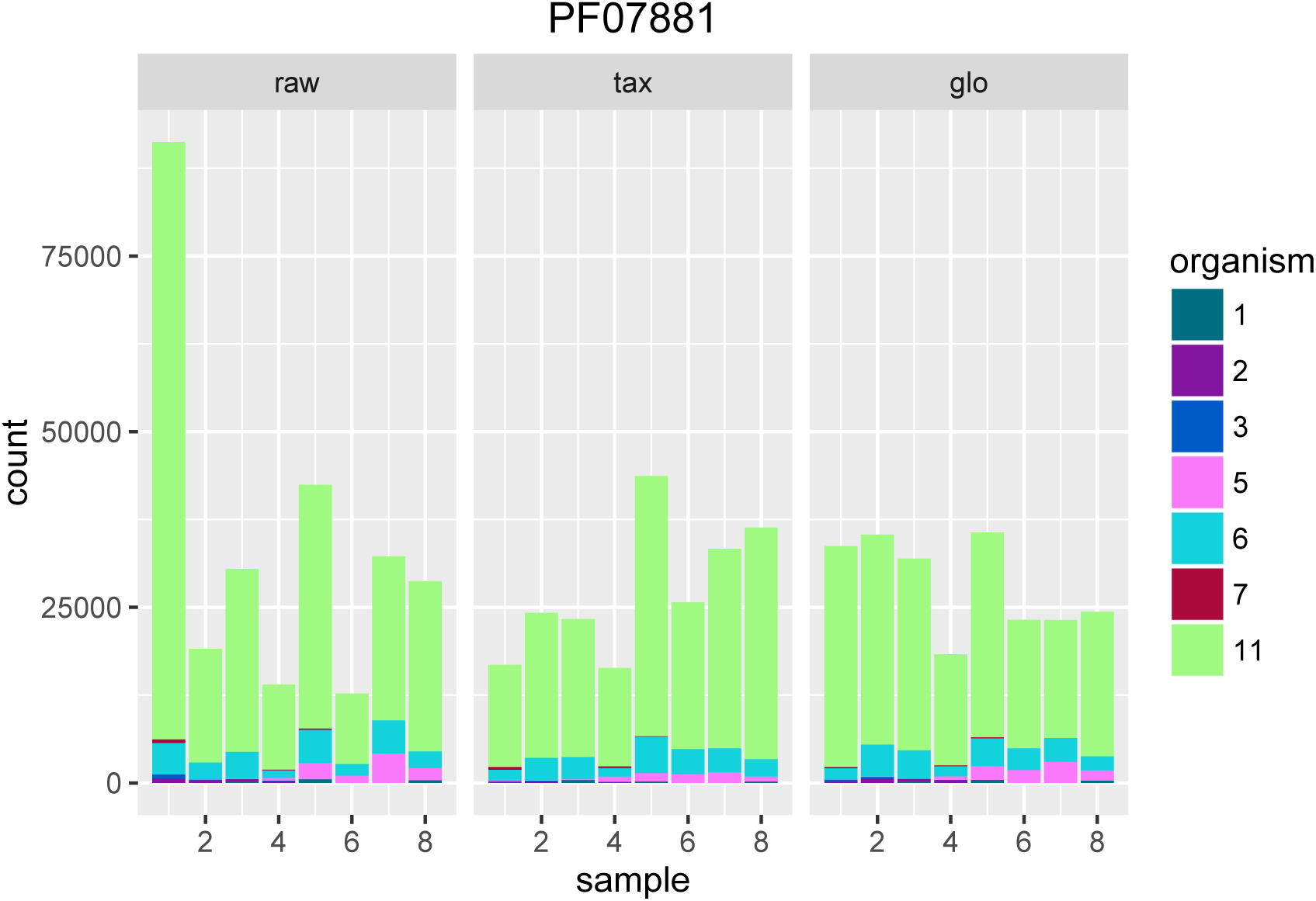
Single feature analysis (PF07881). Stacked bars in three parts (x-axis) show organism-specific counts before scaling (left), after taxon-specific scaling (middle) and after global scaling (right). Loss of significance due to (mis)scaling of profiles from mainly one organism. Feature is significant for organisms Org2, Org5 and Org11 in transcriptome analysis. Taxon-specific and global scaling result in adjusted p-values 1.79e^−3^ and 0.66, respectively.

**Figure 13.**
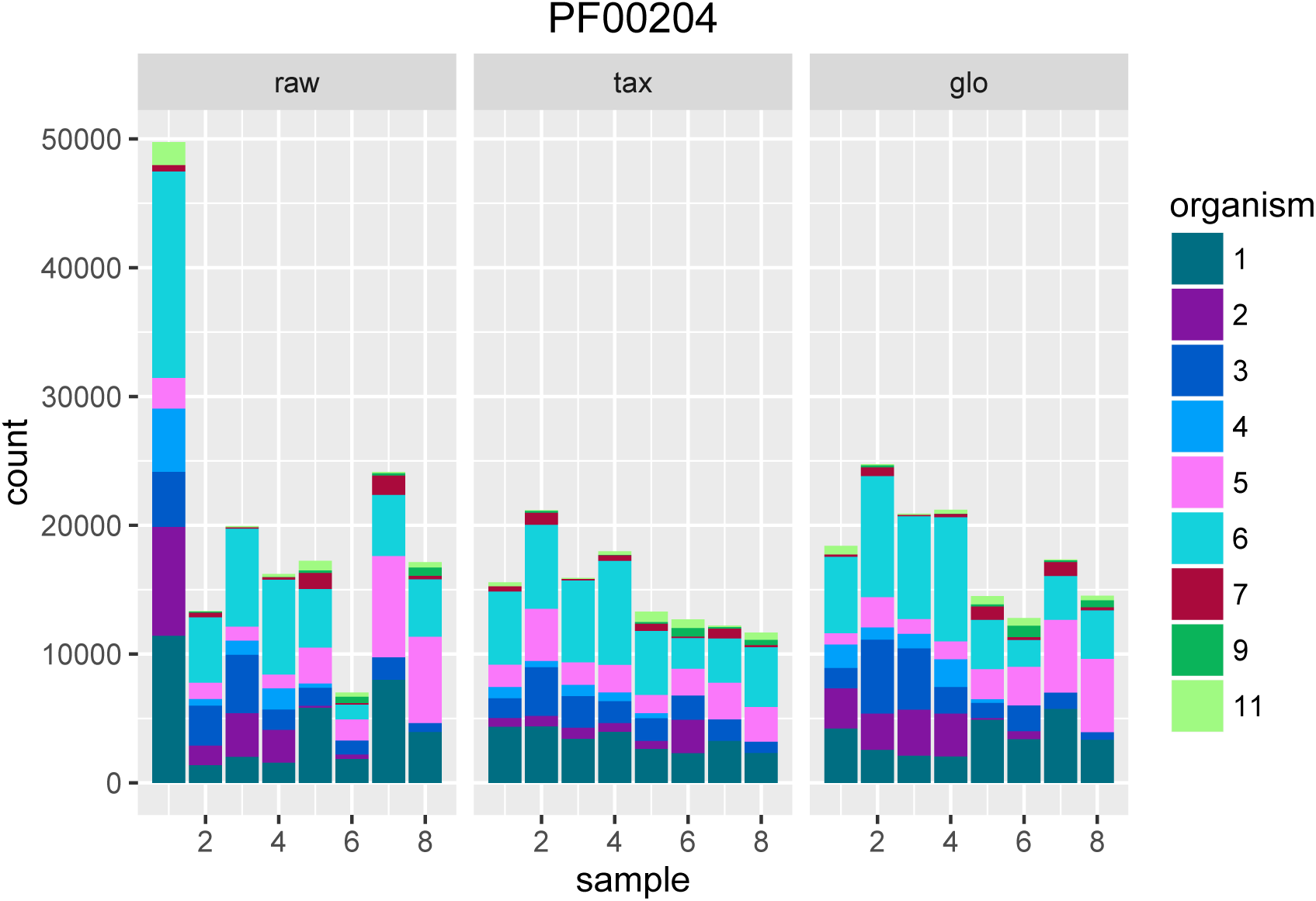
Single feature analysis (PF00204). Stacked bars in three parts (x-axis) show organism-specific counts before scaling (left), after taxon-specific scaling (middle) and after global scaling (right). With boosting effect observable for both methods, i.e. feature is insignificant in transcriptome analyses and significant for metatranscriptomic counts. For Org1 and Org6 the adjusted p-values are close to 0.05 for single analysis, with taxon-specific scaling an adjusted p-value *p* = 0.005 is achieved.

The prevalent effect for both normalization methods was cancellation. Often, the DEF from multiple transcriptomes cancel each other out (see Fig. 14 and Additional File 2: Fig. 13) and in some cases an organism switch could be observed (see Additional File 2: Fig. 14 and 15).

**Figure 14.**
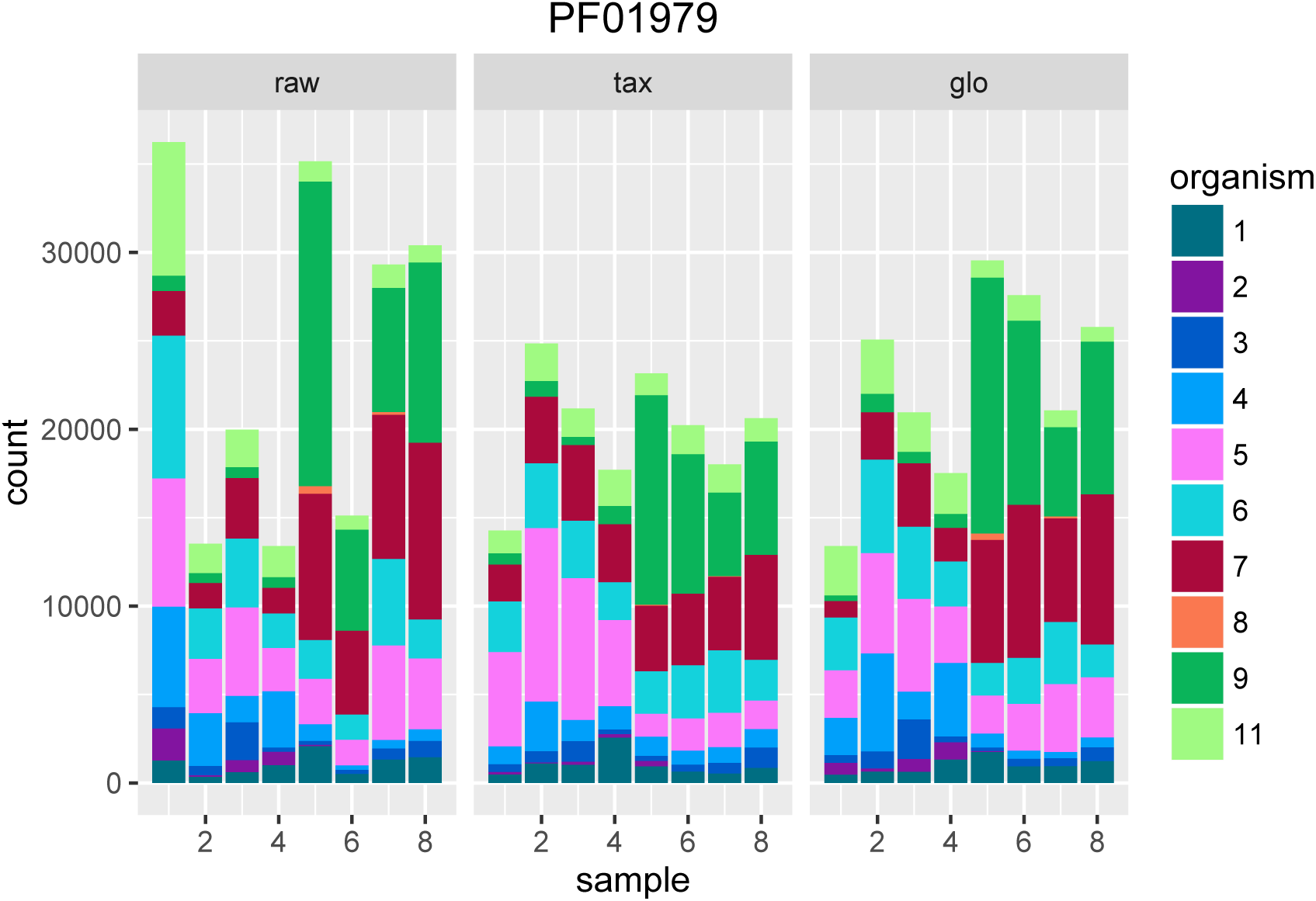
Single feature analysis (PF01979). Stacked bars in three parts (x-axis) show organism-specific counts before scaling (left), after taxon-specific scaling (middle) and after global scaling (right). Feature is significant for transcriptome analysis but becomes insignificant for metatranscriptomic counts (cancellation effect), i.e. the adjusted p-value with both scaling methods is above 0.05.

**Figure 15.**
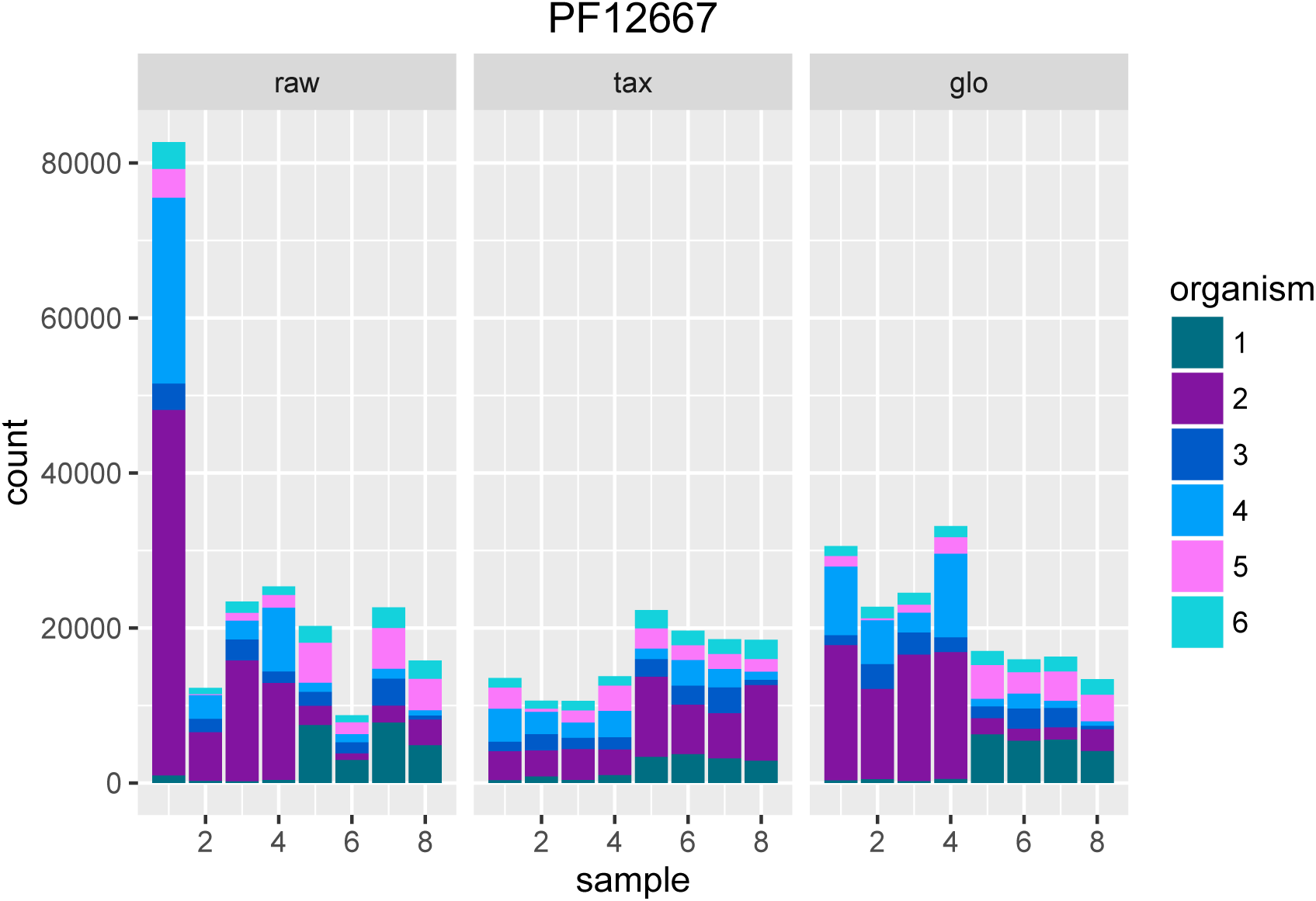
Single feature analysis (PF12667). Stacked bars in three parts (x-axis) show organism-specific counts before scaling (left), after taxon-specific scaling (middle) and after global scaling (right). Significant feature with opposite direction for the two scaling methods. Taxon-specific and global scaling result in adjusted p-values 6.91e^−5^ and 1.33e^−5^, respectively. The log2 fold change for condition A in comparison to condition B is 0.70 for taxon-specific scaling and -0.82 for global scaling.

An example for a contradicting expression direction is shown in Fig 15. For this feature, the incorrect scaling factors obtained by global scaling for Org2 and Org4 suggest a higher expression of this feature in “day 13”, while taxon-specific scaling predicted the expression to be higher in the “day 27” condition.

## CONCLUSIONS

Differential expression analysis in metatranscriptomics is challenging. Metatranscriptomic count data from RNA-Seq experiments show two main modes of biological variation. The functional composition of transcripts reflects the activity of organisms and systematic changes might indicate a metabolic response to experimental conditions. The taxonomic composition of transcripts can change as well and a change may not necessarily be explainable in terms of controlled experimental conditions. In contrast to metagenomics, in metatranscriptomics the questions ”who is there?” and ”what are they doing” are not necessarily connected and need to be answered separately. If the two questions are not separated, there is a considerable risk, to interpret variations in the taxonomic composition as functional changes. This may even happen if the functional profiles of all organisms stay the same under different conditions.

Normalization of metatranscriptomic data must have the goal to eliminate the influence of taxonomic variations from functional analysis. We argue that for a correct normalization the metatranscriptome needs to be decomposed to normalize the organism profiles independently. Then the metatranscriptomic count data may be recombined from the normalized profiles to look for any global trends in the superimposed count data. If differential expression tools are directly applied to the metatranscriptomic count matrix a high risk of erroneous results is encountered. Our simulations indicate that the main risk is not to miss some of the true differences but the real danger is to detect a large number of false functional differences which arise from taxonomic abundance variations across samples. In particular, if these variations are condition dependent the false positive rate can explode, circumventing all statistical control mechanisms for bounding the false discovery rate.

We would like to point out that our findings do not affect metatransciptome studies that just aim to analyze the functional repertoire from RNA-Seq data. The question which functions or genes are expressed is much easier to answer than the question what is the functional response to a change of experimental conditions. However, it is important to note that our results do not only apply to the classic two conditions setup that we used throughout our study. Also for multiple conditions and time series a correct normalization is essential to separate functional from taxonomic trends in the metatranscriptomic composition variations.

## AUTHOR’S CONTRIBUTIONS

HK implemented the method and performed all experiments. PM designed the model and wrote the ”Normalization” section of the manuscript. HK & PM designed the simulations, performed data interpretation and wrote the manuscript. All authors read and approved the final manuscript.

